# Transcriptome Dedifferentiation Observed in Animal Primary Cultures is Essential to Plant Reprogramming

**DOI:** 10.1101/542647

**Authors:** Norichika Ogata

## Abstract

Tissue culture environment liberate cells from ordinary laws of multi-cellular organisms. This liberation enables cells several behaviors, such as growth, dedifferentiation, acquisition of pluripotency, immortalization and reprogramming. Each phenomenon is relating to each other and hardly to determine. Recently, dedifferentiation of animal cell was determined and quantified in increasing information entropy of transcriptome. The increasing information entropy of transcriptome induced by tissue culture may reappear in plant cells too. Here we corroborated it. Measuring information entropies of transcriptomes during reprogramming of plant cells suggested that reprogramming is a combined phenomenon of dedifferentiation and re-differentiation.

## I. INTRODUCTION

Tissue culture is performed to maintain isolated portions of multicellular organisms in an artificial milieu that is outside the individual organism and for considerable periods of time [1]. It is known over a century that cells derived from cultured explants are, in general, different from cells of the corresponding tissue in a living organism [2, 3]. In these tissue cultures, cells are liberated from stimulations and prohibition which is ordinary in multi-cellular organisms [4]. This liberation is essential for growth, dedifferentiation, acquisition of pluripotency, immortalization and reprogramming. However, each phenomena related to each other and some of them had not been scientifically justified, such as dedifferentiation, reprogramming and immortalization. For example, it is not unclear that whether the immortalized cell line has individual cellular immortality or population immortality with gene pool sharing. In other case, historically, proliferations of cultured cells were considered to a result of dedifferentiation [2]. To concrete discussion, concrete definition of each phenomenon which happens in liberated tissue cultures. Recently, the cellular dedifferentiation was quantitive defined as increasing of information entropy of transcriptome [5]. A dedifferentiation of animal cells in primary explant culture was corroborated previously [5]. Then we hypothesized that dedifferentiation of cells in primary explant culture is a common phenomena for diverse multi-cellular organisms. Here we corroborated whether plant cell dedifferentiated in primary explant culture too or not using a shared transcriptome data set [6].

## II. MATERIALS AND METHODS

Transcriptome data set was obtained from DDBJ SRA (DRA000400) [6]. In this entry, time course total RNA sampling during reprogramming of leaf cells of the Moss Physcomitrella patens (0, 1, 3, 6, 12 and 24 hours). Each sample has there biological replicates. We mapped transcriptome sequence data using bowtie 1.1.2 [7] since they used SOLiD sequencer. Information entropy was calculated from all count data as previously described [5]. We compared culture time and information entropy of transcriptome data.

The moss leaf cells were cultured in BCDAT medium. Information entropy of transcriptome was measured in each culture time.

## III. RESULTS AND DISCUSSION

The plant cells dedifferentiated in primary explant culture, equal to animal cells; the information entropy of transcriptome data increased during culture (Figure 1, Figure 2). This result suggested that dedifferentiation of cells in primary explant culture is a common phenomenon for diverse multi-cellular organisms. However, in the case of plants, the information entropy decreases after the information entropy increases up to about 6 hours. This is thought to be re-differentiation to construct new plants and reprogramming can be explained as an integration of dedifferentiation and regeneration. Callus may be regarded as dedifferentiated cells which holding down the re-differentiation process; using callus and plant tissue culture, bi/multi-stability of transcriptome [8, 9] could be demonstrated in plant. If their re-differentiation process is caused by the determination of intercellular division of labor based on cell-to-cell communication [10], it will be possible to examine them in a test that separates cultured cells.

**Fig 1.**
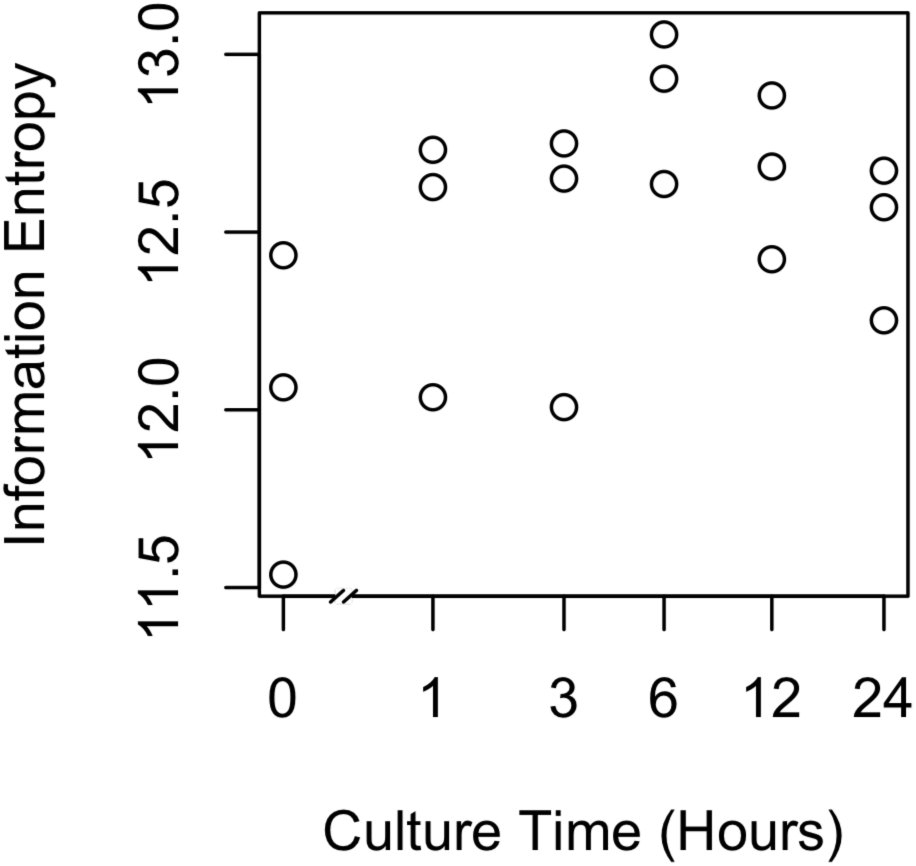
Scatter plot of culture time vs information entropy of transcriptome.

**Fig 2.**
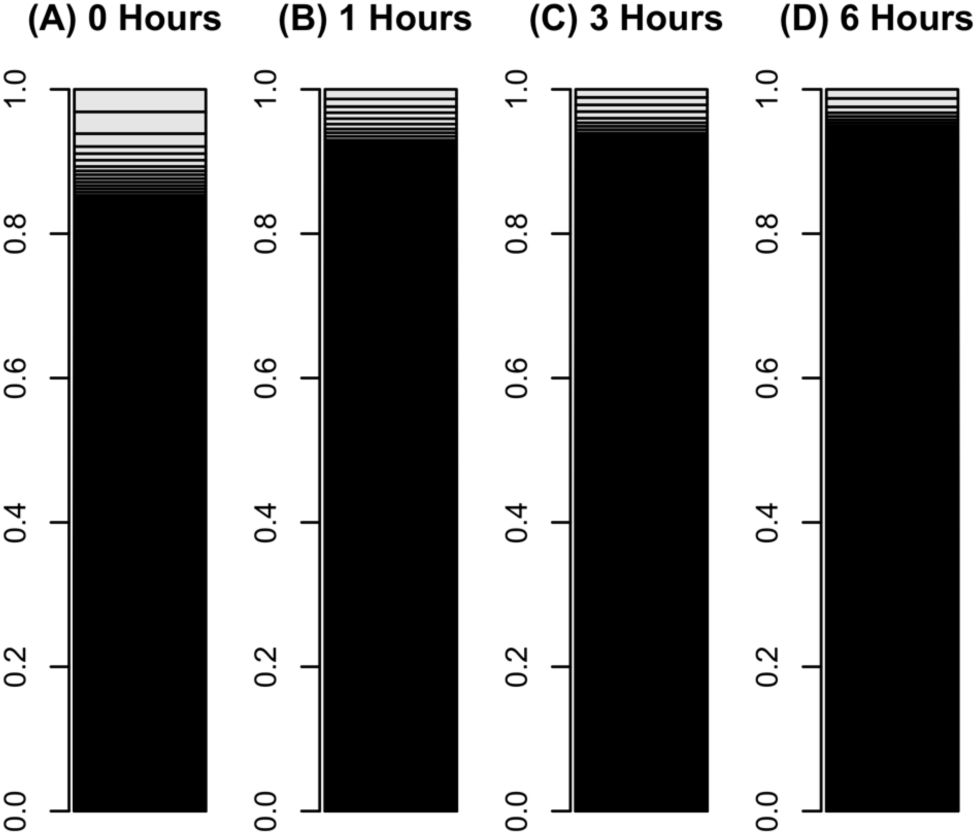
Bar charts of leaf cells transcriptome during reprogramming.

The occupation rate of genes in a transcriptome was plotted in a bar chart. Heights of boxes in a bar chart indicate the occupation rate of genes in a transcriptome. Although more than 50,000 transcripts are included in these bar charts, most are invisible and are included in black regions. (A) A transcriptome of leaf cells cultured for 0 hours in BCDAT medium. (B) A transcriptomes of leaf cells cultured for 1 hour in BCDAT medium. (C) A transcriptomes of leaf cells cultured for 3 hours in BCDAT medium. (D) A transcriptomes of leaf cells cultured for 6 hours in BCDAT medium.

